# HRannot: An accurate and user-friendly gene annotation pipeline for vertebrates

**DOI:** 10.1101/2024.05.21.595213

**Authors:** Siwen Wu, Sisi Yuan, Wei-ming Li, Zhengchang Su

**Affiliations:** Institute of Medical Biology, Chinese Academy of Medical Sciences, Peking Union Medical College, Kunming, Yunnan, China; Department of Bioinformatics and Genomics, the University of North Carolina at Charlotte, Charlotte, NC 28223, USA; The Second Affiliated Hospital of Kunming Medical University, Kunming, Yunnan, China

**Keywords:** Gene annotation, homology-based, RNA-seq data-based, protein-coding genes, pseudogenes

## Abstract

With the development of long reads sequencing technologies and assembly tools, high-quality genome assembly becomes routine in individual labs. In the past few years, numerous high-quality genome assemblies for various vertebrate individuals and species have been deposited in public databases. However, most of these genomes are not annotated for the protein-coding genes, the major working horse in cells, limiting their applications. This dilemma is caused by the reality that the existing easily-used tools cannot achieve accurate annotation, whereas accurate tools such as the NCBI’s eukaryotic genome annotation pipeline are unavailable to individual labs due to the complexity of their use and their high demand for computing resources. Here, we developed an accurate and user-friendly gene annotation pipeline HRannot, enabling accurate annotation of both protein-coding genes and pseudogenes in vertebrate genomes using limited computing resources. Based on both homologous genes in related species and and RNA-seq data from the same species, HRannot is able to annotate both known and novel genes. HRannot outperforms all available well-regarded gene annotation tools, and is comparable or even better in some cases than the NCBI’s gene annotation pipeline.

## Introduction

The Sanger sequencing technique developed in 1977 [1] had been the most commonly used technology to sequence genomes before the advent of next-generation sequencing (NGS) technology [2] developed in 2005. Since 2014, the single molecule real time (SMRT) sequencing technology developed by PacBio Inc. [3] and nanopore-based sequencing technology developed by Oxford Nanopore Technology (ONT) [4, 5] enabled the generation of long reads up to 100 kbp in length. Although the earlier long reads had a high error rate, when combined with highly accurate short reads, they enable high quality chromosome-level genome assemblies. Recent development of highly accurate long reads sequencing technologies and assembly tools has made gap-less telomer to telomer (T2T) genome assembly routine in individual labs [6]. Numerous genomes of different species as well as of different individuals of the same species have been assembled and released by individual labs and large consortia. Particularly, the Vertebrate Genomes Project (VGP) plans to assemble the genomes of all the roughly 70,000 extant vertebrate species, enabling a new era of discovery across the life sciences [7]. The Earth BioGenome Project (EBP) aims to sequence and characterize the genomes of all eukaryotes on Earth over a period of 10 years [8], providing unprecedented opportunities to study the biological process of eukaryotes on Earth. Moreover, hundreds of thousands of individual human genomes have also been sequenced [9-11], and it is expected that more and more genomes will be made available in public databases. However, most of the assembled genomes in public database lack gene annotations. For example, until September 2025, only 6,613 (23%) of 28,754 vertebrate genomes deposited in NCBI’s GenBank have the annotation for at least one coding sequence (CDS) [12].

Clearly, accurate annotation of all genes and pseudogenes in the genome of an organism is important to understand the biology and evolutionary history of the organism. And in fact, many gene annotation tools have been developed, and they can be roughly classified into *de novo* -based methods, RNA-based methods, and homology-based methods. Tools combining multiple methods have also been developed [13-15]. However, none of the existing tools could sufficiently and accurately annotate both protein-coding genes and pseudogenes. *De novo* gene annotation tools such as AUGUSTUS [16], GENSCAN [17] and GeneMark-ES [18] tend to have high false-positive and false-negative rates. RNA-based tools such as Trinity [19] are often limited to genes expressed in the sampled tissues or experimental conditions. Homology-based tools such as Genomix [20], GeMoMa [21] and Scipio [22] are likely to miss new genes that are unique to the reference genome. Combined tools such as Braker3 [13] (combining AUGUSTUS [16], GeneMark-ETP [23] and RNA-seq reads mapping), CAARs [14] (combing GeneMark-ETP [23] and RNA-seq reads mapping) and FINDER [15] (combing gene family alignments and RNA-seq reads mapping) tend to have high false-positive rates. Although NCBI [24] and ENSEMBL [25] have developed eukaryotic genome annotation pipelines and made great efforts to generate high-quality annotations for the so-called reference genome of each sequenced species, these tools are not run in a fully automatic mode, and often need tedious manual corrections. In addition, these pipelines are not available to individual research groups.

To fill these gaps, we developed an accurate and user-friendly gene annotation pipeline named HRannot. By using a combination of homology-based and RNA-seq data-based approaches, HRannot is able to annotate both protein-coding genes and pseudogenes in a broad spectrum of vertebrates accurately. By comparing with the state-of-the-art gene annotation tools, we find that HRannot outperforms them and is comparable or even better in some cases than the NCBI’s annotation pipeline. Thus, HRannot is a useful tool for annotate genes and pseudogenes in an increasing number of assembled genomes.

## Materials and Methods

### Overall flowchart of HRannot

We designed HRannot to predict both known and novel genes in a genome by combining a homology-based method and an RNA-seq-based method. Moreover, HRannot is also designed to annotate both protein-coding genes and pseudogenes in the genome. As shown in Figure 1, both the homology-based method and RNA-seq-based method consist of a data preparation procedure and a gene prediction procedure.

**Figure 1.**
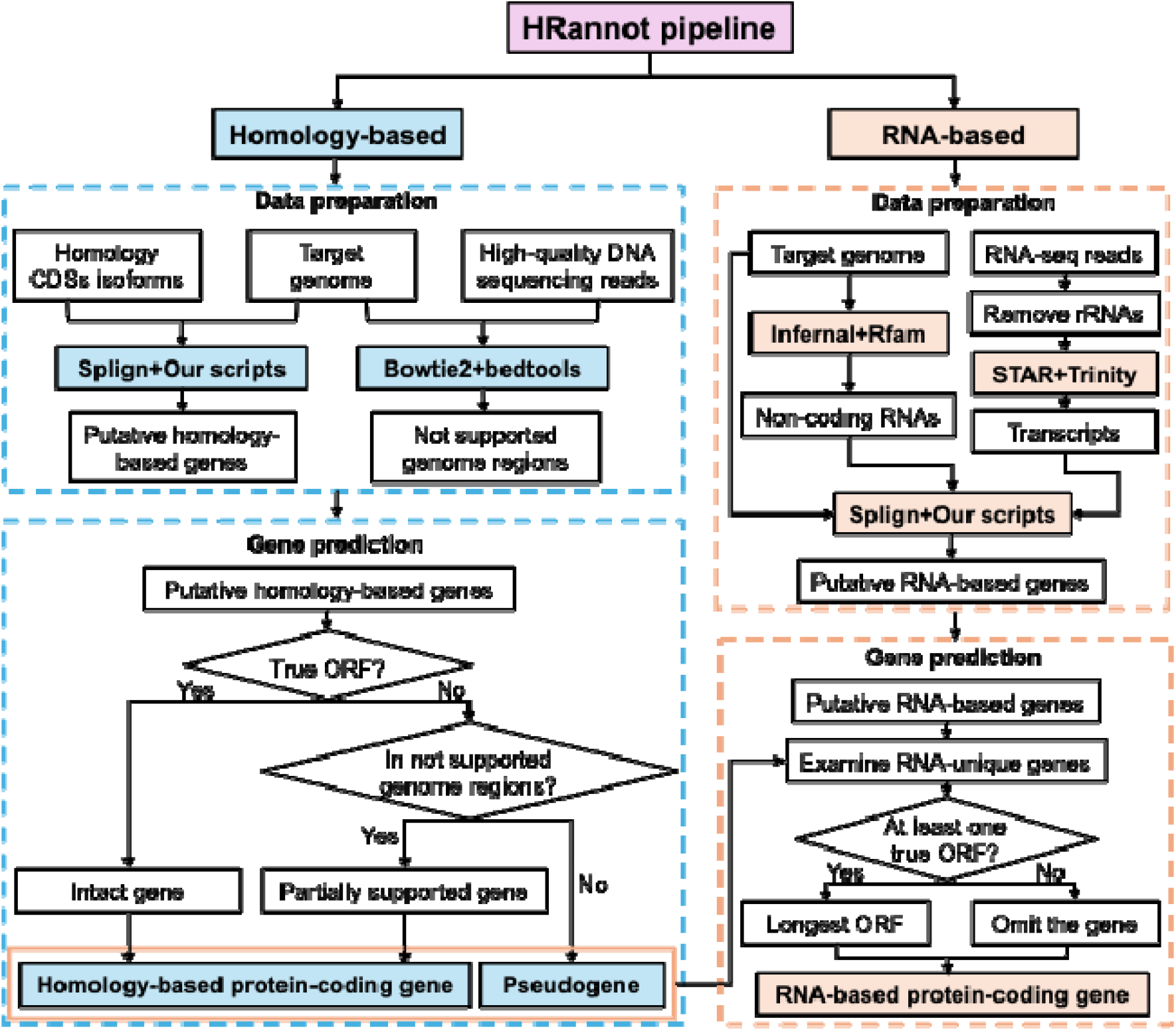
Flowchart of the HRannot pipeline. HRannot predicts protein-coding genes using a combination of homology-based and RNA-based approaches. After proper data preparations, HRannot first uses the homology-based method to predict homologous genes based on each gene model in each reference genome in an order described in the main text, and then it employs the RNA-based method to predict genes that are missed by the homology-based method using RNA-seq data.

### The Homology-based method

In the data preparation step, given a target genome to be annotated, we collected a set of reference genomes with gene annotations. Ideally, the set consists of as many as possible well-annotated genomes in the same class as the target genome. We map the CDSs of each genes in each reference genome to the target genome using Splign (2.0.0) with default settings [26]. We sort the reference genomes in the descending order according to their average CDS mapping identity to the target genome. Moreover, we map the high-quality DNA sequencing reads (Illumina DNA short reads or HiFi long reads) from the same individual using bowtie (2.4.1) [27] with no gaps and mismatches permitted.

In the prediction step, for each reference genome, for each of its annotated genes whose CDSs can be mapped to the same contig/scaffold/chromosome of the target genome, we predict each mapped region in the target genome to be an exon/CDS and the intervals between the putative exons/CDS to be introns. We concatenate all the putative CDSs to form a putative full-length CDS, and predict the corresponding locus (including the putative CDSs and introns) to be an intact genes, if and only if the putative full-length CDS forms an open reading frame (ORF), i.e., it has a length of an integer time of three and its last three nucleotides form the only stop codon. If the putative full-length CDS contains a premature stop codon (nonsense mutation), or its length is not an integer time of three (ORF shift mutation), we check whether the locus can be completely covered by at least n (by default, n=10, or n= ¼ of the average genome coverages) DNA short reads from the same individual. If the answer is yes, we consider the pseudogenization (nonsense mutation or ORF shift mutation) is supported by the high-quality DNA sequencing reads, and predict the sequence to be a pseudogene. Otherwise, we consider the pseudogenization is not supported by the high-quality sequencing reads, and call the sequence a partially supported gene, because the pseudogenization might be artificially caused by errors of the assembled target genome. We call these intact genes, partially supported genes, and pseudogenes to be the predictions based on the reference genome.

To combine all the predictions based on different reference genomes, we initiate the current predictions as those based on the first genome in the set of sorted reference genomes, and update them iteratively by considering each of the predictions based on each remaining reference genomes in their resorted order. More specifically, for each putative gene/pseudogene *g* based on the next reference genome, we replace with a gene/pseudogene q in the current predictions, if *g* overlaps *q* by more than 0.95 of q’s length (the number could be set by users) and *g*’s template mapping identity is higher than that of *q*’s. If *g* does not overlap any of the genes/pseudogenes in the current predictions, and its template mapping identity is greater than 0.99 (the number could be set by users), we added *g* to the current predictions. We consider all intact genes and partially supported genes as homology-based protein-coding genes (Figure 1).

### RNA-seq-based method

In the data preparation step, we first collect as many as possible RNA-seq data from various tissues of the target species or its closely related species. We then map all the RNA-seq reads to the rRNA database [28] using bowtie (2.4.1) with default settings [27]. We remove the reads that map to rRNAs.

In the prediction step, we first predict non-coding RNA genes using infernal (1.1.2) [29] with Rfam as the reference database [30]. We then align the cleaned reads to the target genome using STAR (2.7.0c) [31] with default settings and assemble transcripts using Trinity (2.8.5) [19] with its genome-guided option. We next map the well-assembled transcripts to the target genome using Splign (2.0.0) [26] with default settings, and remove those that at least partially overlap non-coding RNAs, protein-coding genes or pseudogenes predicted by the homology-based method. For each of the remaining mappings, we use the same approach mentioned in the homology-based method to obtain the putative full-length CDS. If the putative full-length CDS contains at least one ORF, we predict the longest one to be a protein-coding gene if it is at least 300 bp (the number could be determined by users) and has an average identity of at least 0.985 (the number could be determined by users) (Figure 1).

### Annotations of the human, mouse, chicken, Tibetan ground-tit and turtle genomes

To evaluate HRannot and other state-of-the-art genome annotation tools, we downloaded the best-assembled genomes available in GenBank for humans (T2T-CHM13v2.0), mice (PAHARI_EIJ_v1.1), chickens (GRCg6a), Tibetan ground-tits (PseHum1.0) and turtles (GSC_CCare_1.0) with accession numbers GCF_009914755.1, GCF_900095145.1, GCF_000002315.6, GCF_000331425.1, and GCF_023653815.2, respectively. We downloaded high-quality DNA sequencing data collected from the host individual of each genome from the GEO database with the accession numbers listed in Table S1. For homology-based annotations, we selected a set of reference genomes consisting of all annotated genomes of species in GenBank’s RefSeq, belonging to the same taxonomic class as the target species. More specifically, these include 242 mammalian genomes (Table S2) for the human and mouse genomes, 145 avian genomes (Table S3) for the chicken and Tibetan ground-tit genomes, and 47 reptile genomes (Table S4) for the turtle genome. Gene annotation files of these genomes as well as those of the target genomes are downloaded from GenBank using corresponding accession numbers (Tables S1-S4). For RNA-based annotations, we downloaded RNA-seq data sets for each species from the GEO databases, whose accession numbers are listed in Table S1. For computational efficiency, we only used RNA-seq data from the individual or cell line whose assembled genome would be annotated, except for the chicken genome, for which we used 40 datasets collected from 10 tissues of four individual chickens [32].

### Comparison of HRannot with state-of-the-art methods AUGUSTUS, Braker3, GeMoMa, and the NCBI annotation pipeline

For AUGUSTUS [16], we used its *de novo* function with default settings. For Braker3 [13], we used its ETP mode (annotate using both RNA-seq data and protein data) with default settings. For GeMoMa [21], we used both RNA-seq data and homologous annotations with default settings. For the homology-based predictions of GeMoMa [21], we used the same reference genomes as used in HRannot for each target genome (Tables S2-S4).

### Assessment of genes overlapping with NCBI’s annotations

We define a predicted gene/pseudogene *q* to match an NCBI annotated gene/pseudogene *g* if *q* overlaps more than 50% length of g and the identity of the mapping parts are 100%.

### BUSCO assessment

We used BUSCO (5.7.1) [33] to assess the quality of the annotated CDSs and exons of the protein-coding genes and pseudogenes in each species by each tool. More specifically, for the human and mouse genomes, we used mammalia_odb10; for the chicken and Tibetan ground tit genomes, we used aves_odb10; and for the turtle genome, we used tetrapoda_odb10.

### Precision and Recall assessment

For the precision and recall of each tool against NCBI’s annotations in each genome, we used the following formulas:

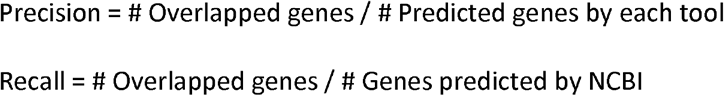

## Results

### The homology-based and RNA-based predictions of HRannot are complementary with each other

HRannot starts with the homology-based approach to predict homologous genes in a target genome using annotated CDSs in multiple reference genomes, and then uses the RNA-seq-based approach to predict genes that are missed by the former approach (Materials and Methods). To evaluate HRannot, we applied it with default settings to the best-assembled genomes in GenBank for five vertebrate species including humans, mice, chickens, Tibetan ground-tits and turtles (as of October 2025), using the default settings with the selected sets of reference genomes and RNA-seq datasets for each species (Materials and Methods). To see the relationship between the number of predicted homology-supported genes and the number of reference genomes used, we plot the accumulative number of predicted genes in each target genome as a function of the number of reference genomes with the highest mean mapping identities used (Materials and Methods). As shown in Figures 2a-2e, the few reference genomes with the highest mean mapping identities are able to predict the vast majority of homology-supported genes in each target genome, and the accumulative number of predicted genes saturates after more than 20 reference genomes with the highest mean mapping identities are used. This is particularly true in the human, mouse, chicken and Tibetan ground-tit genomes compared to the case of the turtle genome, due probably to the better annotations of reference genomes in mammals and birds than those of reptiles. These results suggest that the reference genomes that we selected for each target genome are adequate for homology-based gene predictions, and that using more than 20 reference genomes with the highest mean mapping identities contributes little to the total number of predicted homology-supported genes. The number of predicted homology-supported protein-coding genes and pseudogenes using all the selected reference genomes in each species are listed in Tables 1 and 2. Most of these genes are covered by RNA-seq reads from the tissues of each species, suggesting that they are transcribed and thus are likely to be authentic.

**Table 1:** Comparison of the annotations of different tools from several aspects.

| Species | Category | HRannot |  | AUGUSTUS |  | Braker3 |  | GeMoMa |  | NCBI |  |
| --- | --- | --- | --- | --- | --- | --- | --- | --- | --- | --- | --- |
|  |  | Protein-coding gene | Pseudogene | Protein-coding gene | Pseudogene | Protein-coding gene | Pseudogene | Protein-coding gene | Pseudogene | Protein-coding gene | Pseudogene |
| Human | # Annotation by the tool | 20717 | 784 | 23489 | - | 24869 | - | 59002 | - | 20090 | 15534 |
|  | # Overlap with NCBI's annotation | 15340 | 134 | 15424 | - | 11425 | - | 13456 | - | - | - |
|  | BUSCO completeness | 91.1% | - | 61.3% | - | 81.4% | - | 91.2% | - | 99.3% | - |
| Chicken | # Annotation by the tool | 18272 | 541 | 15898 | - | 18984 | - | 29054 | - | 17485 | 240 |
|  | # Overlap with NCBI's annotation | 13071 | 9 | 12370 | - | 12080 | - | 13300 | - | - | - |
|  | BUSCO completeness | 96.7% | - | 71.9% | - | 92.1% | - | 97.7% | - | 97.7% | - |
| Mouse | # Annotation by the tool | 20624 | 56 | 10897 | - | 18848 | - | 39398 | - | 20482 | 5091 |
|  | # Overlap with NCBI's annotation | 15667 | 24 | 9116 | - | 13361 | - | 15385 | - | - | - |
|  | BUSCO completeness | 93.2% | - | 35.5% | - | 86.0% | - | 92.5% | - | 98.4% | - |
| Turtle | # Annotation by the tool | 19670 | 1952 | 19641 | - | 20848 | - | 38766 | - | 20259 | 705 |
|  | # Overlap with NCBI's annotation | 14862 | 170 | 15955 | - | 12956 | - | 13309 | - | - | - |
|  | BUSCO completeness | 94.7% | - | 53.7% | - | 89.7% | - | 90.9% | - | 97.8% | - |

**Table 2:** Comparison of gene annotation results of the Tibetan ground-tit genomes by HRannot and NCBI’s pipeline.

| Methods | Homology-based |  |  | # RNA-based genes | # Total protein-coding genes | # Total pseudogenes | BUSCO completeness |
| --- | --- | --- | --- | --- | --- | --- | --- |
|  | # Intact genes | # Partially supported genes | # Pseudogenes |  |  |  |  |
| HRannot | 15,352 | 448 | 81 | 102 | 15,902 | 81 | 99.0% |
| NCBI | - | - | - | - | 15,814 | 83 | 99.5% |
| Common | - | - | - | - | 14,030 | 0 | - |

To see the relationship between the predicted RNA-supported genes and the RNA-seq datasets used, we choose datasets from 1, 2, …, 10 chicken tissues and used them to predict RNA-supported genes. As shown in Figure 2f, the accumulative number of predicted RNA-supported genes increases rapidly with the increase in the number of datasets used, followed by a slow linear increase, suggesting that using additional diverse datasets can potentially predict more RNA-supported genes. The number of predicted RNA-supported protein-coding genes using all the selected RNA-seq datasets in each species are listed in Tables 1 and 2. Notably, the number of RNA-supported genes are much smaller than that of homology-supported genes in each species, as the RNA-based approach is designed to predict genes that are missed by the homology-based method (Materials and Methods).

**Figure 2.**
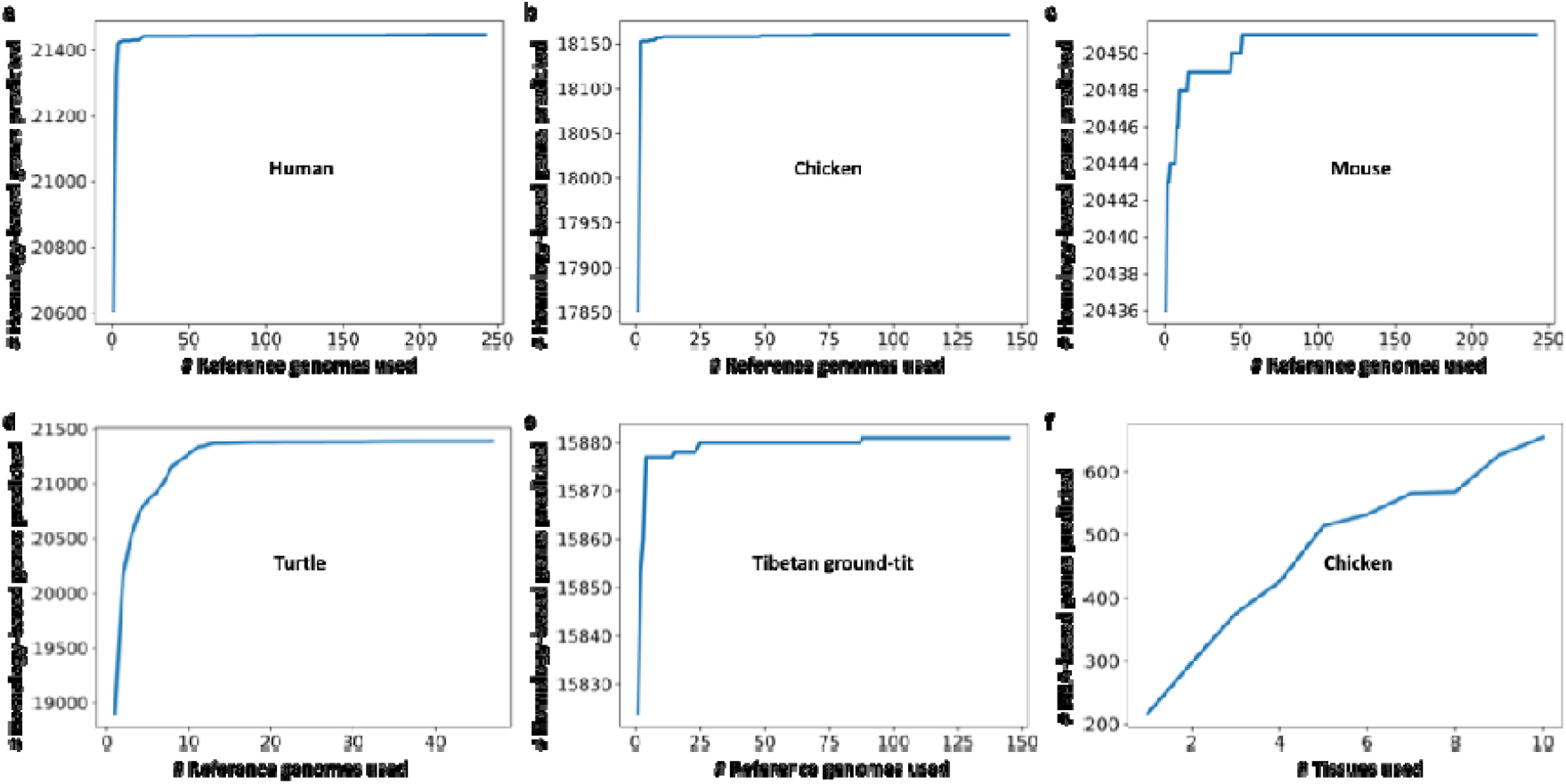
Relationship between the number of predicted homologues genes/RNA-based genes and the number of reference genomes/tissues whose RNA-seq data are used. **a**. The accumulative number of predicted homologues genes in the human genome as a function of the number of reference mammalian genomes used. **b**. The accumulative number of predicted homologues genes in the chicken genome as a function of the number of reference bird genomes used. **c**. The accumulative number of predicted homologues genes in the mouse genome as a function of the number of reference mammalian genomes used. **d**. The accumulative number of predicted homologues genes in the turtle genome as a function of the number of reference reptile genomes used. **e**. The accumulative number of predicted homologues genes in the Tibetan ground-tit genome as a function of the number of reference bird genomes used. **f**. The accumulative number of predicted RNA-based genes in the chicken genome as a function of the number of tissues whose RNA-seq data are used.

### HRannot outperforms the state-of-the-art gene annotation tools in the human, mouse, chicken and turtle genomes

We compared the results of HRannot in four target genomes (human, mouse, chicken and turtle) with those of three state-of-the-art tools AUGUSTUS [16], Braker3 [13] and GeMoMa [21], obtained by applying each of them to the four target genomes using the default settings of the tool and the same sets of reference genomes and RNA-seq datasets if the tool support such datasets as inputs. We chose AUGUSTUS [16], Braker3 [13] and GeMoMa [21] for comparison, as they represent *de novo*-based, RNA-seq + protein-based, and homology + RNA-seq-based annotation methods, respectively. As NCBI has made great efforts to annotate these genomes, we used the NCBI’s annotations as the standard for evaluations. We first compared the number of genes predicted by each tool with those annotated by NCBI. As shown in Table 1, for all the four species, HRannot achieved the closest number of protein-coding genes (20,717, 18,272, 20,624 and 19,670, for human, chicken, mouse and turtle, respectively) to those of NCBI’s annotations (20,090, 17,485, 20,482 and 20,259, for human, chicken, mouse, turtle, respectively). By contrast, GeMoMa [21] predicted the largest number of protein-coding genes for each species (59,002, 29,054, 39,398 and 38,766 for human, chicken, mouse, turtle, respectively), which is 1.7-2.9 times higher than those of NCBI’s annotations (Table 1). Both AUGUSTUS [16] and Braker3 [13] predicted considerably fewer or more genes than NCBI’s annotation pipeline (Table 1).

Next, we compared the tools for their ability to recall the genes annotated in each genome by NCBI. As shown in Table 1, in the human and turtle genomes, AUGUSTUS [16] recalled the highest numbers of NCBI-annotated genes (15,424 and 15,955, respectively), while HRannot achieved the second highest numbers (15,340 and 14,862, respectively). In the chicken genome, GeMoMa [21] recalled the highest number (13,300) and HRannot had the second highest number (13,071), while GeMoMa [21] predicted 10,782 more genes than HRannot (Table 1). In the mouse genome, HRannot recalled the highest number (15,667), and GeMoMa [21] had the second highest number (15,385) (Table 1). Moreover, AUGUSTUS [16] tends to have higher precision with the trade-off of lower recall; and braker3 [13] and GeMoMa [21] tends to have higher recall with the trade-off of lower precision (Table 1, Figure 3). In contrast, HRannot always had the most balanced performance for precision and recall among the tools (Table 1, Figure 3).

**Figure 3.**
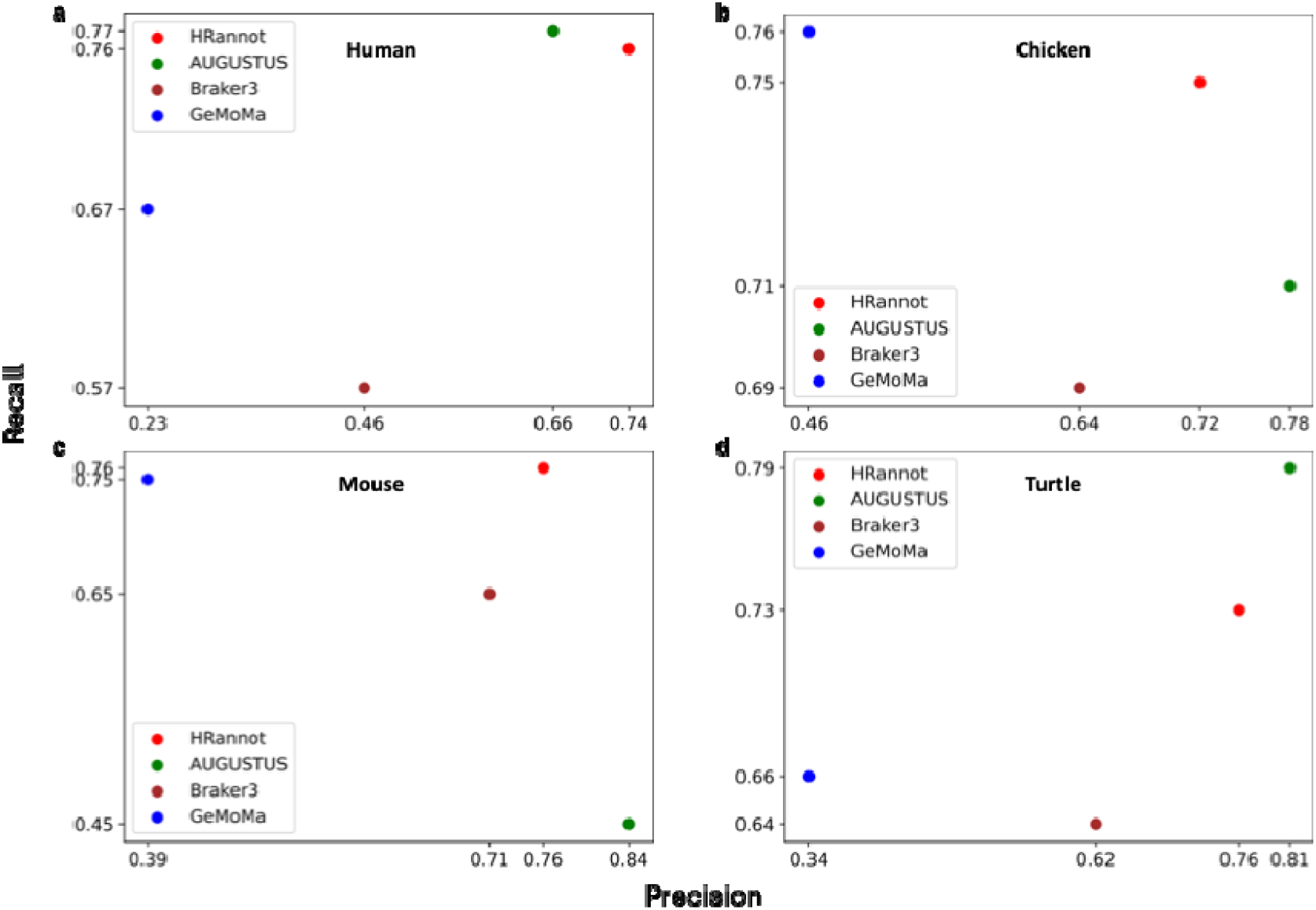
Precision and Recall assessment of the annotations by different tools for the human (a), chicken (b), mouse (c) and turtle (d) genomes.

Finally, we compared the tools by the BUSCO [33] completeness assessment. HRannot achieved more than 91% completeness in all the four species (Table 1), suggesting that its annotations of these genomes are quite complete. Although GeMoMa [21] obtained similarly high BUSCO [33] completeness values (90.9%-97.7%) in the genomes, however, it produced too many false positives (Table 1). Braker3 [13] had BUSCO [33] completeness values arranging from 81.4% to 92.1% in the genomes, which is much lower than those obtained by HRannot and GeMoMa [21] (Table 1). AUGUSTUS [16] obtained the lowest BUSCO [33] completeness values ranging from 35.5% in the mouse genome to 71.9% in the chicken genome (Table 1), suggesting that it misses a large number of genes in these genomes, although its precision and recall are higher than some other tools, because it generates many fragmental predictions (Figure 4, Table S5). More specifically, HRannot achieved the highest BUSCO [33] completeness value in mouse and turtle (Table 1). In human, HRannot was only 0.1% lower than that of GeMoMa [21], while GeMoMa [21] predicted 38,285 more protein-coding genes than HRannot (Table 1). In chicken, GeMoMa [21] achieved 1% higher BUSCO completeness value than HRannot, while GeMoMa predicted 10,782 more protein-coding genes than HRannot (Table 1). By checking the details of the BUSCO [33] assessment results, we found that except for chicken, HRannot had the highest value of complete and single-copy BUSCOs (Figure 4, Table S5), suggesting that there is little over-annotation. The complete and duplicated BUSCOs value is lower than 1% for HRannot in all the four annotations (Figure 4, Table S5), suggesting again that HRannot achieves a complete but few redundant annotations.

**Figure 4.**
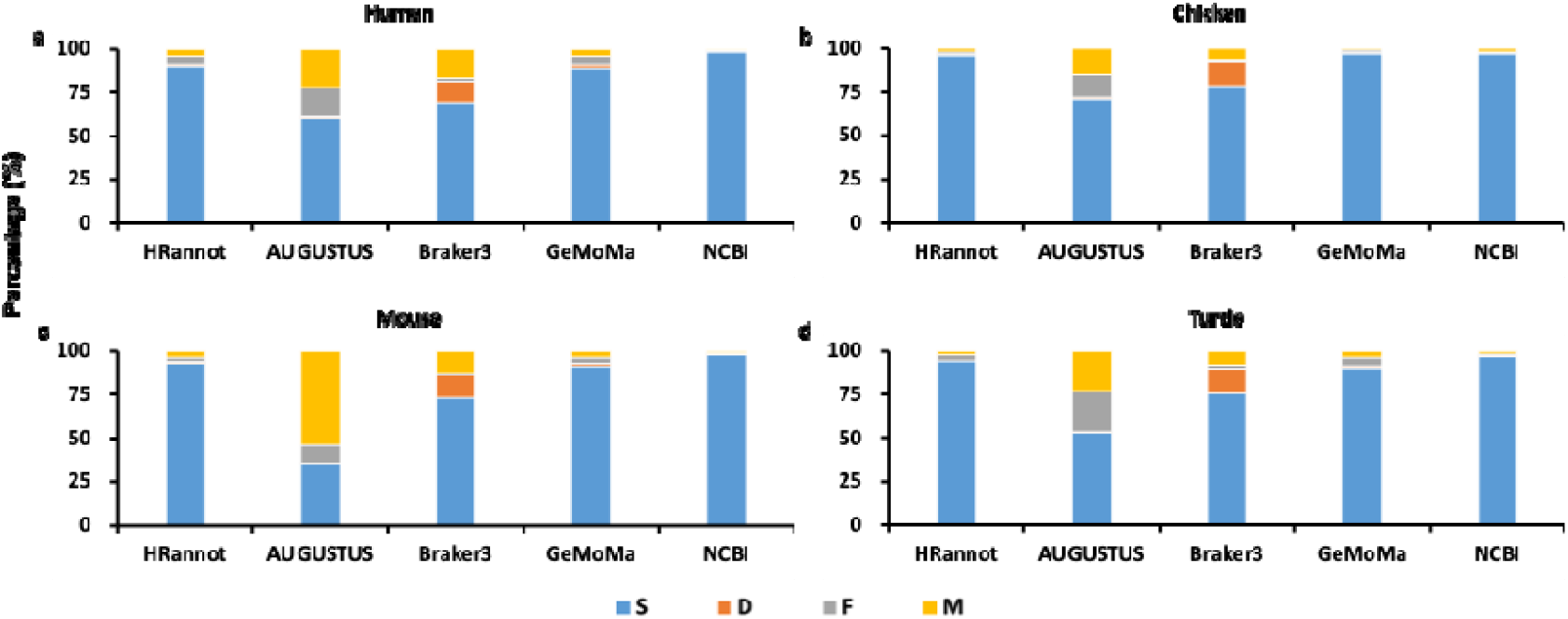
BUSCO completeness of the annotations in different genomes. **a-d**. BUSCO completeness values of the annotations from different annotation tools for human, chicken, mouse and turtle genomes, respectively. In each subfigure, S represents Complete and single-copy BUSCOs, D represents Complete and duplicated BUSCOs, F represents Fragmented BUSCOs, and M represents Missing BUSCOs. Specifically, BUSCOs completeness in Tables mentioned above consists of S and D.

In addition, all the other three tools do not annotate pseudogenes in the target genomes. In contrast, as shown in Table 1, HRannot predicted 784, 541, 56 and 1,952 pseudogenes in the human, chicken, mouse and turtle genomes, respectively, with 134, 9, 24 and 170 overlapping with NCBI-annotated pseudogenes in the four genomes, respectively. Notably, all the pseudogenes predicted by HRannot were supported by the high-quality sequencing reads of the corresponding individual of the species (Methods and Materials), suggesting that the pseudogenes predicted by HRannot are reliable. However, since HRannot mainly predicts unprocessed pseudogenes, and NCBI mainly predicted processed pseudogenes, there are a few overlaps between the pseudogenes predicted by the two tools.

### HRannot are comparable or even outperforms the NCBI annotation pipeline on the Tibetan ground tit genome

Although HRannot outperforms the three state-of-the-art methods in almost all the measures computed, it underforms the NCBI annotation pipeline due probably to the fact that greater efforts have been made by the NCBI team to annotate these genomes using larger volumes of RNA-seq data available in these species, particularly, the human, mouse and chicken genomes. To further demonstrate the prediction accuracy of HRannot, we used it to annotate the genome of a Tibetan ground-tit individual (Table S1), a less-studied avian species with limited availability of RNA-seq data, and compared the results with the annotations from NCBI. By using the CDS isoforms of 145 well-annotated avian species from NCBI RefSeq (Table S3) as the references, the homology-based method of HRannot predicted 15,352 intact genes, 448 partially supported genes and 81 pseudogenes (Table 2). By using the RNA-seq reads from multiple tissues of the individual (Table S1), the RNA-based method of HRannot annotated 102 genes for the individual (Table 2). In total, HRannot predicted 15,902 protein-coding genes and 81 pseudogenes for the genome (Table 2), while NCBI annotated 15,814 protein-coding genes and 83 pseudogenes in it (Table 2). There are 14,030 protein-coding genes shared by the two pipelines, suggesting that HRannot is able to annotate most of the protein-coding genes (88.7%) predicted by the NCBI annotation pipeline (Table 2). HRannot achieved a similar BUSCO completeness value (99.0%) to that of NCBI’s annotation (99.5%) (Table 2). However, there is no overlap of the pseudogenes predicted by the two pipelines (Table 2).

To find the reason for the inconsistent protein-coding genes and pseudogenes predicted by the two pipelines, we examined the gene-bodies of the unique genes and pseudogenes predicted by each method. We found that NCBI’s pipeline could mistakenly predict our predicted protein-coding genes as pseudogenes, or mistakenly predict our predicted pseudogenes as protein-coding genes. For examples, genes *PTCD2* and *DIO3* are annotated as protein-coding genes by the NCBI’s pipeline, while they are annotated as pseudogenes by HRannot since there are stop codons in the middle of their ORFs (Figures 5a and 5b); gene *LOC*102109485 (named *LOC*107212260 in our annotation) is annotated as a pseudogene by the NCBI’s pipeline, while it is annotated as protein-coding genes by HRannot since there are neither stop codons nor ORF shift mutations in its ORFs (Figure 5c). Thus, these results suggest that our HRannot is more accurate in annotating some genes and pseudogenes.

**Figure 5.**
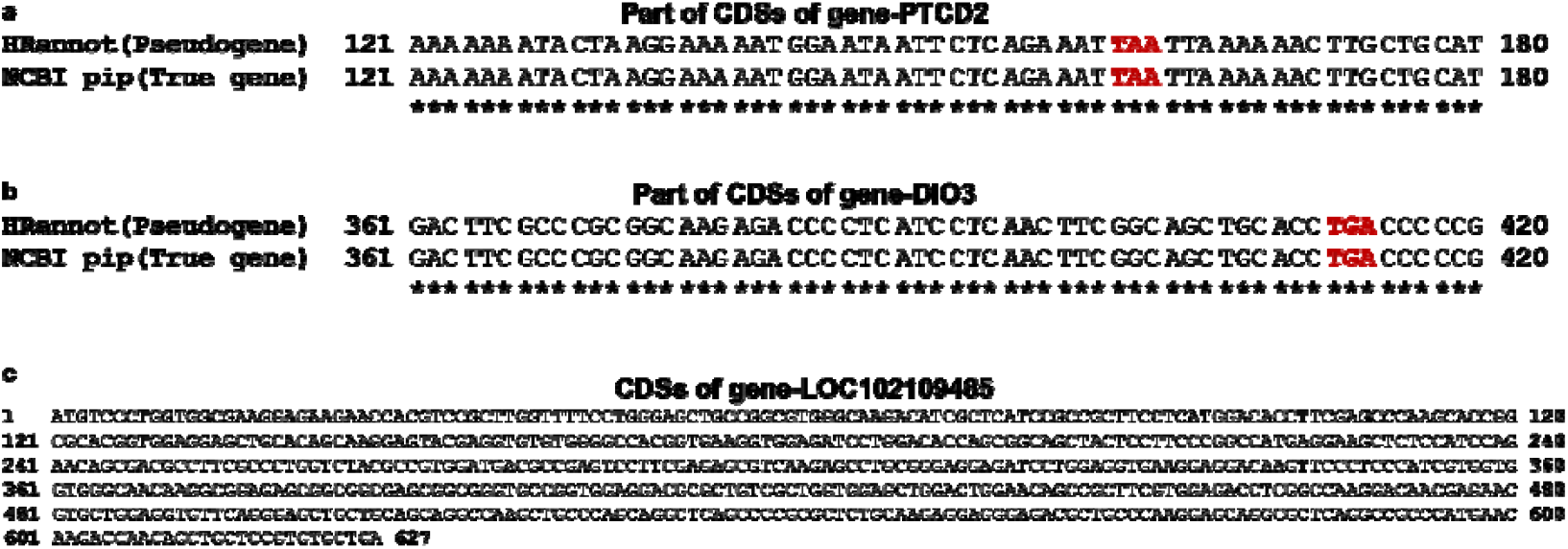
Examples of different annotations for the same gene/pseudogene from the NCBI and HRannot annotation pipelines. **a**. Part of ORFs of gene *PTCD2* annotated by both the two pipelines contains a stop codon labeled in red. **b**. Part of ORFs of gene *DIO*3 annotated by both the two pipelines contains a stop codon labeled in red. **c**. The ORF of gene *LOC*102109485 annotated by the two pipelines. The sequence contains no pseudogenization mutations.

## Discussion

We developed a homology- and RNA-based gene annotation pipeline HRannot that demonstrates superior performance compared to several established tools, including AUGUSTUS [16], Braker3 [13] and GeMoMa [21], three popular and widely used gene annotation tools. Through comprehensive benchmarking, our pipeline consistently achieved higher annotation accuracy, as evidenced by a BUSCO completeness score [33] exceeding 91% across diverse test vertebrate genomes (Table 1). Importantly, our method maintains a well-balanced trade-off between precision and recall, addressing a common limitation observed in other annotation tools that often sacrifice one for the other (Figure 2). While Braker3 [13] and GeMoMa [21] rely heavily on homology and transcript evidence, respectively, our pipeline integrates both sources more effectively, resulting in improved gene model prediction. Compared to AUGUSTUS [16], which primarily depends on de novo prediction, our pipeline shows substantial gains in both structural accuracy and biological relevance. These results underscore the robustness and reliability of our approach, making it particularly valuable for annotating newly sequenced or non-model genomes.

Besides, HRannot is also comparable or even more sensitive in some cases compared with the NCBI gene annotation pipeline, i.e. correcting some pseudogenes or protein-coding genes that are mistakenly annotated as the other type by the NCBI’s pipeline (Figure 5). In addition, a particular advantage of our pipeline lies in its ability to identify unprocessed pseudogenes, which retain intron-exon structures and closely resemble their functional counterparts. In contrast, the NCBI’s pipeline primarily annotates processed pseudogenes, which lack introns and often arise from retrotransposition. The detection of unprocessed pseudogenes is especially challenging due to their structural similarity to active genes and their typically low expression levels. Our pipeline’s success in capturing these elements highlights its enhanced sensitivity and specificity in distinguishing subtle genomic features. Although the BUSCO [33] completeness value of NCBI’s annotations are always higher than those of HRannot, there are always many manual corrections of NCBI’s annotation, while HRannot is completely automatic, causing the relatively lower BUSCO [33] completeness value.

Given its demonstrated effectiveness, our pipeline HRannot offers a compelling alternative to current standards and holds promise for broad applications in genomics research.

## Declarations

### Consent for publication

Not applicable.

### Availability of data and materials

The annotation results of the five species using HRannot are available at https://doi.org/10.6084/m9.figshare.30462191.v1. The code and pipeline description for HRannot are available at https://github.com/zhengchangsulab/HRannot.

## Authors’ contributions

ZS supervised and conceived the project; SW and ZS designed the pipeline; SW, SY, WL and ZS performed data analysis; and SW and ZS wrote the manuscript.

